# Semantic coding in the occipital cortex of early blind individuals

**DOI:** 10.1101/539437

**Authors:** Sami Abboud, Denis A. Engemann, Laurent Cohen

**Affiliations:** Institut du Cerveau et de la Moelle épinière, ICM, Inserm U 1127, CNRS UMR 7225, Sorbonne Université, F-75013, Paris, France; Parietal project-team, INRIA, Universite Paris-Saclay, 91120, Saclay, France; Service de Neurologie 1, Hôpital de la Pitié Salpêtrière, AP-HP, F-75013, Paris, France

**Keywords:** Plasticity, visual system, blindness, semantics, language, MEG, MVPA

## Abstract

The visual cortex of early blind individuals is reorganized to support cognitive functions distinct from vision. Research suggests that one such prominent function is language. However, it is unknown whether the visual cortex of blind individuals codes for word meaning. We addressed this question by comparing neuronal activity evoked by a semantic decision task, using magnetoencephalography (MEG), between 12 early blind and 14 sighted participants otherwise comparable with regard to gender, age and education. We found that average brain responses to thousands of auditory word stimuli followed similar time courses in blind and sighted participants. However, in blind participants only, we found a sustained enhancement of activity in the visual cortex. Moreover, across the whole brain, we found an effect of semantic category from about 400 ms after word onset. Strikingly, in blind participants, semantic categories were discriminable starting 580 ms after word onset from signal captured by sensors sensitive to the visual cortex. We replicated the analyses in time windows locked to stimulus onset and behavioral response, using both classical hypothesis testing and machine learning for single-trial classification. Semantic decisions were well classified in all participants (AUC ∼ 0.60), but generalization capacity across participants was found reduced in the blind group due to a larger variability of discriminative patterns. In conclusion, our findings suggest that brain plasticity reorganizes the semantic system of blind individuals, and extends semantic computation into the visual cortex.

## Introduction

Blindness from birth abolishes visual input into the occipital cortex and initiates a chain of events that lead to its involvement in various non-visual functions (reviewed in Ricciardi and Pietrini, 2011; Voss and Zatorre, 2012; Bedny, 2017; Abboud and Cohen, 2019). Those include language processing, a high-order cognitive functions which recruits the occipital cortex of early blind individuals. Both general language components such as speech comprehension, word generation and verbal memory, and more selective features such as sentence-level structure and syntactic complexity were found to activate the occipital cortex (e.g. Amedi *et al.*, 2003; Bedny *et al.*, 2011; Lane *et al.*, 2015).

Along those lines, there is evidence that the occipital cortex in blind individuals also plays a role in semantic processing. First, visual regions as early as primary visual cortex (V1) are activated when contrasting heard words and pseudowords (Bedny *et al.*, 2011). Second, stimulating the occipital pole using repetitive transcranial magnetic stimulation leads to semantic errors during verb generation (Amedi *et al.*, 2004). Third, a contrast between a task involving the semantic content of words and a task of speaker gender identification on reversed words, activates the left fusiform, middle occipital and superior occipital gyri (Noppeney *et al.*, 2003). Finally, occipital activity in the gamma-band (75-110 Hz) is sensitive to semantic congruency between sounds and tactile objects (Schepers *et al.*, 2012).

However, showing occipital activation during a verbal semantic task versus baseline, or finding differential activation to conditions requiring different degrees of semantic processing, does not amount to demonstrating the existence of actual coding of word meaning in those regions. Actually, there is evidence that blind and sighted subjects share the same overall organization of their semantic system, covering the lateral and ventral temporal cortices, the inferior parietal cortex and the prefrontal cortex (Huth *et al.*, 2016; Ralph *et al.*, 2017). The activation profile of the left middle temporal gyrus during semantic similarity judgments on verb pairs is similar across blind and sighted participants (Bedny *et al.*, 2012). Moreover, in a task involving semantic content, when comparing words referring to hand actions (e.g. tapping) to words referring to sound (e.g. siren), vision (e.g. flash) and motion (e.g. gallop), both blind and sighted participants activated the left posterior middle temporal cortex. Also, when comparing words referring to vision to other categories of words, both blind and sighted participants activated the left inferior temporal gyrus (Noppeney *et al.*, 2003). A similar picture also emerges for the organization of the ventral occipitotemporal cortex according to semantic domains, an organization which persists in blindness (Mahon *et al.*, 2009; Peelen *et al.*, 2013; Wang *et al.*, 2015; van den Hurk *et al.*, 2017). For example, when comparing animals to manmade objects during a size-judgment task, the lateral preference for animals and medial preference for objects prevails in both blind and sighted participants (Mahon *et al.*, 2009). However, such evidence, while highlighting strong commonalities, falls short of demonstrating that blind individuals possess a semantic system identical to that of the sighted, untouched by the large-scale plasticity implicating their occipital cortex in language processing.

We addressed this question by probing the spatiotemporal unfolding of semantic access in early blind participants with high temporal granularity. To this end, we acquired magnetoencephalography (MEG) recordings in blind and sighted participants while they performed a semantic decision task to heard nouns belonging to three semantic categories: animals, plants and manmade objects. Using univariate and multivariate analyses on the sensor array and in source-space, we tested whether and when semantic categories could be discriminated in single participants. We then compared semantic discrimination between blind and sighted participants to identify systematic group differences in semantic processing. Finally, we tested whether the neural implementation of semantic processing is similar across participants and across groups.

## Materials and Methods

### Participants

The study included 12 early blind individuals that were either born blind or have lost their sight during the first six weeks of life (7 males; age: 46±16, mean±SD) and 14 sighted controls (7 males; age: 47±15), out of which 12 were matched in sex, age and education level to the blind group. All participants were native French speakers. None of the blind subject had any form of light perception (Table 1). All participants signed an informed consent form, were paid for their participation and were naive about the aims of the study. The study was approved by the local ethical committee.

**Table 1.**
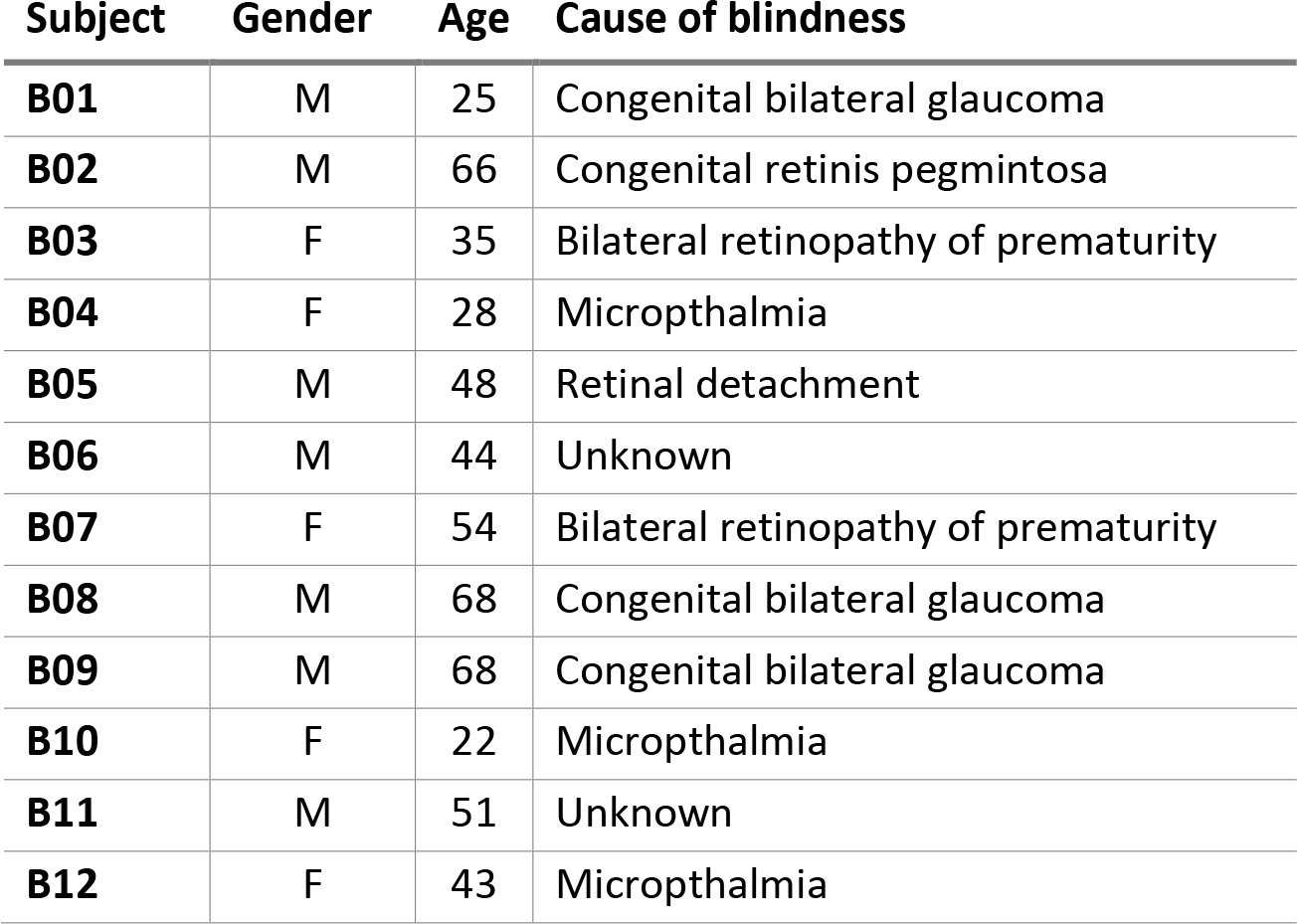
Causes of blindness.

| **Subject** | **Gender** | **Age** | **Cause of blindness** |
| --- | --- | --- | --- |
| **B01** | M | 25 | Congenital bilateral glaucoma |
| **B02** | M | 66 | Congenital retinis pegmintosa |
| **B03** | F | 35 | Bilateral retinopathy of prematurity |
| **B04** | F | 28 | Microphthalmia |
| **B05** | M | 48 | Retinal detachment |
| **B06** | M | 44 | Unknown |
| **B07** | F | 54 | Bilateral retinopathy of prematurity |
| **B08** | M | 68 | Congenital bilateral glaucoma |
| **B09** | M | 68 | Congenital bilateral glaucoma |
| **B10** | F | 22 | Microphthalmia |
| **B11** | M | 51 | Unknown |
| **B12** | F | 43 | Microphthalmia |

### Experimental design and Stimuli

Participants performed an auditory semantic decision task while Magnetoencephalography (MEG) was concomitantly recorded (Fig. 1). Stimuli consisted of words from three semantic categories: Animals, Manmade objects and Plants (see Supplementary Table 1 for a full list). The task was divided into 36 blocks. During each block participants categorized words belonging to a category pair. All three possible pairs were used in 12 blocks. Each block started with two presentations of instructions, mapping word categories to left and right response buttons (e.g., “Animal, Left. Plant, Right”). All three word-categories were equally often associated with left and right responses. Instructions were followed by 3 seconds of silence, and then by a list of 60 words with a uniformly jittered inter-stimulus interval (ISI) of 1.3-1.55 s. Participants were instructed to respond to each word using their left and right index fingers, as fast as they could without sacrificing accuracy. Once the ISI had passed, the next word was presented regardless of the subject’s response. Each block was followed by 5 seconds of rest. The experiment was divided in 12 runs of approximately 7 minutes. Participants were offered a break every three runs but were granted more or less pauses between runs depending on their reported state of fatigue.

**Figure 1.**
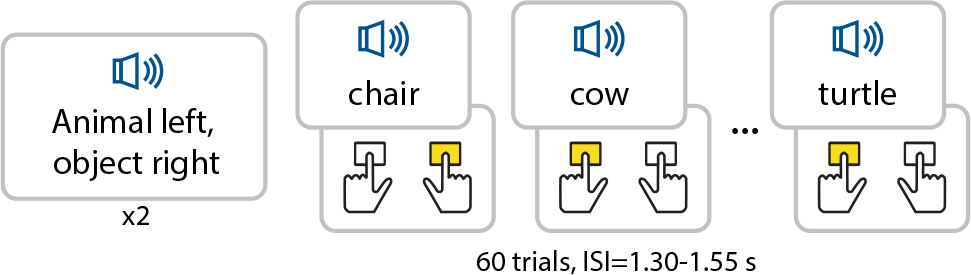
Semantic decision task. Participants were first presented with two instances of instructions indicating the mapping between categories and response buttons (e.g. Animal, left – Object, right). Then, participants had to categorize 60 auditory words per block. Button mapping was balanced across category pairs and the latter were balanced across blocks. Inter stimulus interval was jittered between 1.30 and 1.55 s.

The stimuli totaled to 90 unique words (Lexique database; New *et al.*, 2004), 30 per semantic category (Supplementary Table 1). Words were also equated across three phonemic categories as per their first vocalic phoneme. Those phonemes were the French [a], [e] and [o]. Each semantic category, then, contained ten words from each phonemic category. Auditory stimuli were normalized with regard to relevant statistical parameters and read by the computer (see section *experimental design and stimuli* in Supplementary material).

For each subject and each category, we computed accuracy and median latency, which were analyzed using a mixed-effects ANOVA with category as within-subject factor, group as between-subject factor and participants as random factor. Trials with responses faster than 200 ms or slower than 1400 ms, and trials with incorrect responses, were discarded from the MEG analysis.

### Procedure

Participants first received a short description of the task followed by a practice block outside the magnetically shielded room. Then, the 90 words were played once in order to ensure that participants were familiar with all of them. For each subject, unfamiliar words were excluded from the analyses (mean number of excluded words: 1.7± std: 2.2, min: 0, max: 7). All participants were blindfolded and kept their eyes closed during the acquisition (except for two blind participants who were not able to voluntarily control their eyelids). Participants underwent a high-resolution anatomical image (T1) acquisition.

### Stimulation

We used Psychophysics Toolbox Version 3 (Brainard, 1997; Kleiner *et al.*, 2007) for MATLAB (Release 2015b; The MathWorks, Inc., Massachusetts, United States) to implement the experimental procedure. Auditory stimulation was delivered using Nicolet TIP-300 (Madison, WI, USA) with Echodia ER3-14A foam ear tips (France). Button presses were recorded using a Cedrus Lumina LSC-400B controller.

### MEG acquisition

MEG signals were acquired using a whole-head MEG system with 102 magnetometers and 204 planar gradiometers (Elekta Neuromag TRIUX MEG system) at a sample rate of 1 KHz and online low-pass filtered at 330 Hz. Electrooculography (EOG) and electrocardiogram (ECG) were simultaneously recorded. ECG electrodes were located at the right clavicle and the lower left quadrant of the abdomen. Vertical EOG electrodes were located above and under the right eye and horizontal EOG electrodes were 2 cm lateral to each eye. The ground electrode was located at the left scapula.

### MEG data processing

To clean the MEG data from environmental artifacts temporal Signal Space Separation (tSSS; Taulu *et al.*, 2005) was performed using the MaxFilter tool (Elekta Neuromag). All remaining data processing steps were performed using the MNE software (Gramfort *et al.*, 2013, 2014). The MEG data was then band-pass filtered 0.1-15Hz with default filter parameter settings. Eye movement and cardiac artifacts were corrected using Independent Components Analysis (ICA). ICA was estimated on raw data across all runs using the FastICA algorithm (Hyvärinen and Oja, 2000). Ocular components were detected using Pearson correlations and cardiac components using cross trial phase statistics (CTPS; Dammers *et al.*, 2008), both, with the MNE-Python default settings.

For source localization, we extracted the anatomical information obtained from individual MRI scans using FreeSurfer (http://surfer.nmr.mgh.harvard.edu/) (see section *anatomical surface reconstruction* in the Supplementary Material for details). To estimate cortical neuronal dynamics from the observed sensor array time series we approximated a numeric solution to the biomagnetic inverse problem using cortically constrained Minimum Norm Estimates (MNE) with L2 regularization, dynamical statistical parametric mapping (dSPM) for noise normalization (Dale *et al.*, 2000) and the samw configuration as in Engemann and Gramfort (2015). For facilitating interpretation, we summarized the inverse solution using the Human Connectome Project (HCP) cortical parcellation (Glasser *et al.*, 2016) with 360 functionally-defined regions of interest (ROI) covering the entire cortical surface. More details can be found under *MEG source localization* in the Supplementary Material.

### Statistical Analysis

#### Sensor Space

We assessed category discrimination within participants and between groups using non-parametric permutation clustering tests (Maris and Oostenveld, 2007). Within participants, we used the F-test for independent samples as contrast function in the non-parametric test. We then contrasted the ensuing sensor-wise F-statistics at the group level using the t-test for independent samples with the permutation-clustering test. Settings were consistent with the values suggested in documentation of the MNE software (Gramfort *et al.*, 2014).

#### Source space

To obtain ROI-wise inference, we then contrasted groups using t-tests for independent samples and obtained inference by comparing the observed t-value against an empirical distribution of t-values under the H0 obtained from 10000 label-wise permutations of the participants between groups (Groppe *et al.*, 2011). Multiple comparisons we adjusted using FDR-control (Benjamini and Hochberg, 1995; Groppe *et al.*, 2011).

#### Decoding

We used a logistic regression classifier with penalized likelihood (L2 norm) to learn how to discriminate single-trial semantic categories from the MEG sensors at a given time point. The regularization parameter fixed at the default value of C = 1. Out-of-sample performance was estimated using k-fold cross-validation within participants (with grouping by words) and leave-one-subject-out cross-validation across participants. All machine learning was performed using the scikit-learn software (Pedregosa *et al.*, 2011). For model inspection, we extracted the learned pattern from the linear model (Haufe *et al.*, 2014) and projected it to the source model using the previously learned minimum norm inverse operator. For details see section *Machine Learning* in Supplementary material. To assess the consistency of decoding success we contrasted our pattern classifier against dummy models that predicted independently from the MEG data solely based on the stratification of class-labels. We computed confidence intervals of continuous decoding performance across participants using the non-parametric percentile bootstrap (Efron and Tibshirani, 1994) over 4000 bootstrap replica obtained from drawing n samples with uniform probability. To obtain dependable p-values, we additionally compared the difference between decoding performance and a dummy classifier and the decoding performance between groups using paired t-tests against an empirically estimated distribution under the null hypothesis using 10000 permutations.

### Data availability

The data used in this paper can be made available upon reasonable request, but because of the sensitive nature of the clinical information concerning the special population studied the ethics protocol does not allow open data sharing.

## Results

### Behavior

Performance in the semantic decision task was high in both groups (Blind: 91.0±6.6%, Sighted: 94.4±4.7%). The main effects of group and category, and the group × category interaction were not significant (Group: F(1,24)=2.57, p=0.12; Category: F(2,24)=2.63, p=0.08; Group × Category F(2,24)=2.08, p=0.14). Average response times stood at 0.875±0.077 s in blind and 0.806±0.108 s in sighted participants. We found a significant main effect of category (F(2,24)=26.58, p<1.0e-5), but no main effect of group nor group × category interaction (Group: F(2,24)=3.45, p=0.08, Group × Category F(2,24)=2.67, p=0.09). A post-hoc Tukey’s test showed that the category effect reflected shorter reaction times for plants compared to animals and objects (mean response times: 0.813, 0.851 and 0.848 s, respectively; Animals – Objects: p=0.748; Animals – Plants: p<1.0e-4; Objects – Plants: p<1.0e-4).

### Neural correlates of the semantic decision task

We were first interested in the global differences between blind and sighted participants when performing the semantic decision task (Figure 1). Therefore, separately for each group, we pooled trials from all categories and computed the average evoked response (Fig. 2A-B). We then estimated the cortical sources of this average evoked-response in each subject and averaged their values inside each ROI of the HCP cortical parcellation atlas (Glasser *et al.*, 2016). We found a significant enhancement of activation in the visual cortex of blind as compared to sighted participants. This effect is already present in some visual areas early after word onset. However, starting 160 ms after word onset, the lateral, medial and inferior aspects of bilateral occipital cortex showed a sustained difference between groups, peaking at 600 ms (P<0.05, FDR-corrected, t-test with non-parametric permutations; Fig. 2C). No ROIs showed a significantly stronger activity in sighted than in blind participants.

**Figure 2.**
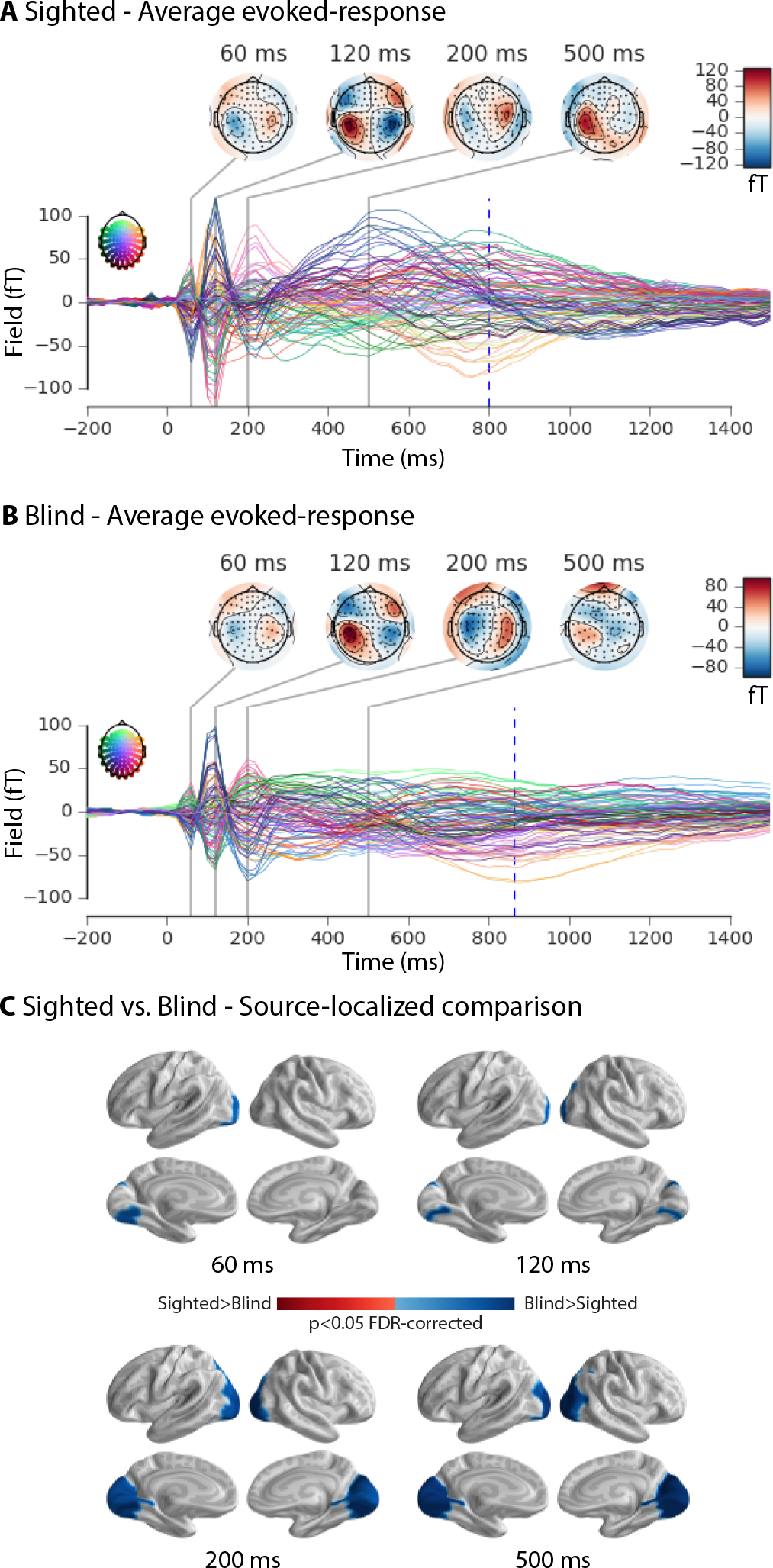
Neural correlates of the semantic decision task. Average evoked-responses in sighted **(A),** and in blind participants **(B)** collapsing across stimuli from all categories. The X-axis shows time (ms) where zero corresponds to word onset and the Y-axis shows the field strength (fT). **(C)** A comparison between sighted and blind participants in source space, showing an enhanced activation in the visual cortex of blind participants (P<0.05, FDR-corrected, t-test with non-parametric permutations). We did not find evidence for enhanced activity in the sighted relative to the blind participants. The results suggest that blind participants additionally recruit the visual system during the semantic decision task.

### Neural signatures of semantic category discrimination

As a first step, we assessed individual category discrimination in successful trials. Exploratory analyses suggested that average MEG signals, in both groups, varied systematically across categories. However, the direction of the effects was highly variable across participants. To summarize category discrimination, we chose an F-statistic with spatiotemporal clustering on sensor data in single participants. This allowed us to generate heat-maps of where differences occurred in time and space while avoiding cancellation when averaging over participants. We found at least one cluster with significant category discrimination in all but one blind and three sighted participants (Fig. 3A). Those clusters overlapped in time starting at about 400 ms for the majority of the sighted and blind participants (Fig. 3B). Spatially, in the majority of the participants of both groups, clusters overlapped over left fronto-central sensors. In the blind, we also observed an overlap of clusters in posterior occipital sensors (Fig. 3C). We then compared individual F-statistics between groups and found one significant cluster where category discrimination was stronger in the blind, encompassing posterior sensors between 580 ms and 900 ms after word onset (P<0.019, t-test with non-parametric permutation clustering; Fig. 3D-E). To delineate the sources showing such group differences in semantic category discrimination, we estimated the cortical sources of the three category-pair contrasts (e.g. animals – manmade) for each subject. We then averaged the source estimates across the three pairs, yielding an individual index of category discrimination, which we compared across groups. Indeed, during the cluster where the F-statistics were stronger in blind participants over posterior sensors, we also found a higher discrimination index in blind participants in sources localized to the lateral, medial and inferior aspects of the occipital cortex, peaking at 740 ms after word onset (Fig. 3F). We then repeated this analysis by aligning epochs on button press rather than on word onset, which we reasoned should increase the sensitivity of the analysis (See section *category discrimination aligned on button press in Supplementary Material)*. The results were indeed equivalent with larger effects (Supplementary Fig. 1A-C) which lead to the adoption of this alignment option in the analyses that follow.

**Figure 3.**
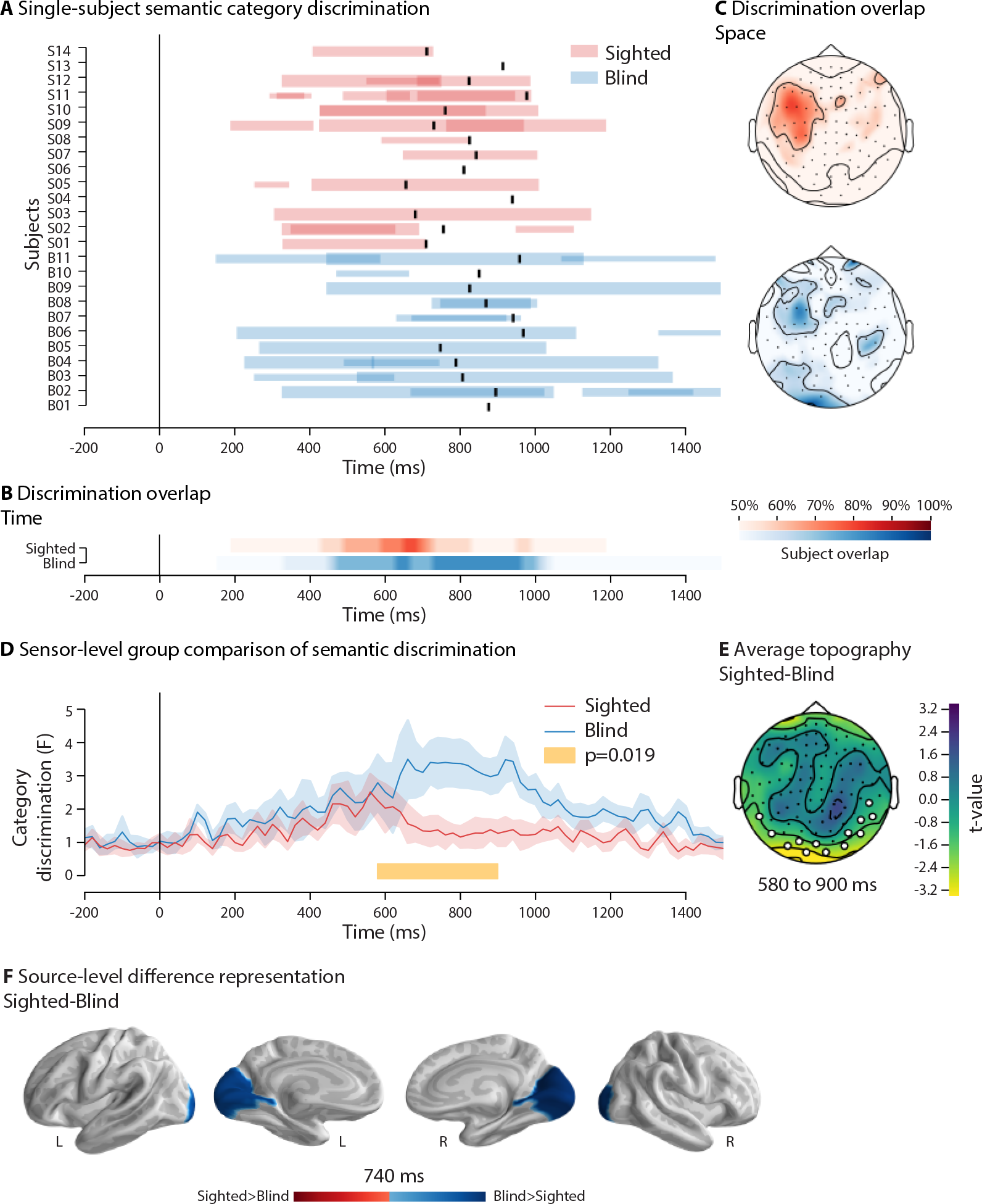
Neural correlates of semantic category discrimination relative to word onset. Panels A-C depict individual analyses. **(A)** The X-axis shows time (ms) where zero corresponds to word onset and each row on the Y-axis shows the result of one subject, sighted in red and blind in blue. Each bar represents a significant spatiotemporal cluster using the magnetometer sensors, bar width is inversely proportional to the p-value, and cluster overlap renders colors darker. At least one significant cluster can be seen in all but one blind and three sighted participants. The black horizontal bars indicate median response times for individual participants **(B)** Temporal overlap of clusters, in sighted (red) and blind (blue) participants. Colors represent the percentage of participants with a significant cluster that includes a given time point. The lightest color represents time points where less than 50% of the participants had a significant cluster. **(C)** Spatial overlap of clusters, showing the percentage of participants with a significant cluster that includes a given sensor. The color scheme follows the description in panel B. Panels D-F depict comparisons between groups. **(D)** The X-axis shows time (ms) where zero corresponds to the word onset and the Y-axis to the average category discrimination in sighted (red) and blind (blue) participants. The yellow bar indicates the temporal extent of the spatiotemporal cluster with significant difference across the groups (P=0.019, spatiotemporal permutation clustering). **(E)** The average topography of the group difference in semantic discrimination during the time of the significant cluster (from 580 to 900 ms). Sensors appearing in the cluster are highlighted in white. **(F)** The peak difference between sighted and blind participants (in the time-window of the cluster in panel D) when comparing the average of all pair-wise contrasts. It shows a stronger effect in the occipital cortex of the blind participants and no regions with a significant effect in the sighted participants. The results suggest that occipital sensors were most sensitive to category discrimination in the blind as compared to the sighted participants and point at specific contributions from the visual cortex.

To further summarize category discrimination, we moved to single-trial decoding using machine learning. This aimed at circumventing individual variability in the spatial layout of semantic responses, and at allowing us to study generalization of semantic coding across words, time, participants and groups. A linear classifier was trained to predict semantic category from MEG signal by learning individual information accessible on the sensor array. We started with sequential decoding, training and evaluating one classifier every 20 ms. In sighted participants, we observed systematic above-chance single-trial classification in the time-window from −480 to +500 ms relative to button press (P<0.05, FDR-corrected, t-test against empirical chance with non-parametric permutation testing; Fig. 4). We also observed significant single-trial classification sparsely at −680, −600, and +580 ms. Classification accuracy reached two local maxima: before button press at −80 ms (AUC=0.581, 95%CI=[0.564, 0.596], P=5.0e-4), and after button press, at +60 ms (AUC=0.594, CI=[0.560, 0.628], P=5.0e-4). In blind participants, we found continuously significant single-trial classification in the time-window from −580 to +540 ms (P<0.05, FDR-corrected; Fig. 4). In addition, we also observed above-chance classification sparsely at −800 to −720, −680 to −640, +640, and +940 ms. As in the sighted, we observed two local maxima of classification accuracy in blind participants: before button press at −160 ms (AUC=0.587, CI=[0.567, 0.603], P<0.0011), and after button press at +80 ms (AUC=0.614, CI=[0.583, 0.645], P<0.0011). Comparing classification accuracy between groups did not yield significant effects (P=0.178; FDR-corrected). This indicates that the temporal architecture of semantic processing is equivalent in sighted and blind participants. As response times were shorter for plants than for the other categories, we wished to verify that the success in decoding between two categories did not depend on response time differences. To this end, we computed bivariate Pearson correlations to probe whether individual decoding scores were correlated with absolute response time differences. This was done using the decoding scores and response time differences of all subjects in all three category pairs. We found negative, weak, non-significant trends in both groups (Blind: r=−0.15, P=0.366; Sighted: r=−0.18, P=0.283), providing no suggestion of a link between decoding success and reaction time differences.

**Figure 4.**
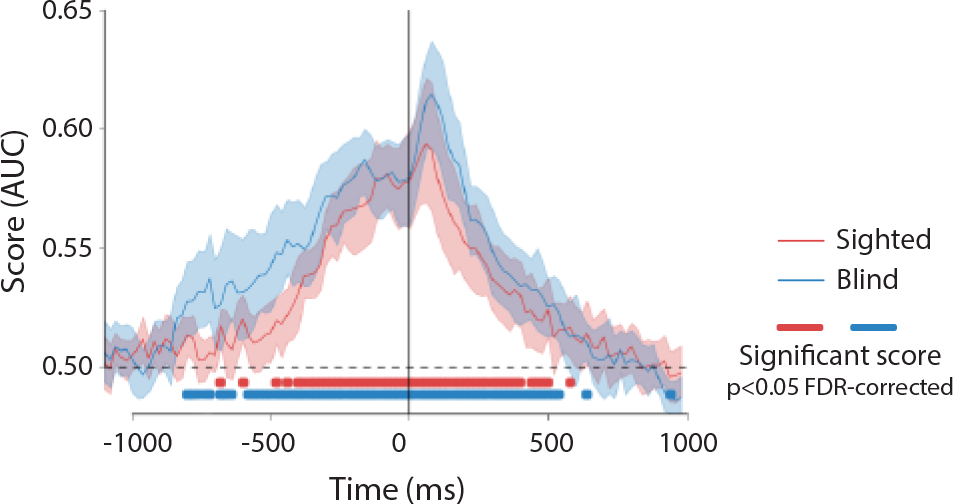
Single-trial decoding of semantic category discrimination. Classification accuracy scores in the sighted (red) and the blind (blue). Scores reflect the group average of single-subject results across the 3 category pairs. X-axis shows time and (ms) where zero corresponds to the button-press and Y-axis shows the AUC score. Shaded areas indicate bootstrapped 95% confidence intervals. Time-points marked with red (blue) indicate statistically significant prediction scores when compared to empirical chance levels in sighted participants (blind; P<0.05, FDR-corrected, t-test with non-parametric permutations). No significant differences were found between the groups. The results suggest that the temporal organization of category discrimination was virtually identical in the sighted and blind participants.

To further assess the contribution of the occipital lobes, we then identified the brain regions which provided the classifier with critical semantic information, by estimating the cortical sources of the neuromagnetic patterns derived from the coefficients of the model underlying the MVPA (Haufe *et al.*, 2014). During the entire time period when both groups showed continuous significant classification (i.e. from −480 ms to +500 ms), the main sources common to both groups were the anterior and lateral temporal lobes and the left-predominant inferior frontal cortex (Fig. 5a). We compared the groups over this entire time window, finding the visual cortex to be significantly more influential in the blind than in the sighted (P<0.05, FDR-corrected; t-test with non-parametric permutations; see Fig. 5b for an illustration at the classification peak in the blind). Conversely, the prefrontal cortex was marginally more influential in the sighted than in the blind participants (P<0.05, uncorrected; Fig. 5b). Results at −80, +60 and +80 ms (i.e. the three other classification maxima, two in the sighted and one in the blind), all showed a similar pattern (Supplementary Fig.s 2 and 3). Taken together, these results converge with the univariate analysis and provide evidence for semantic category coding in the visual cortex of the blind.

**Figure 5.**
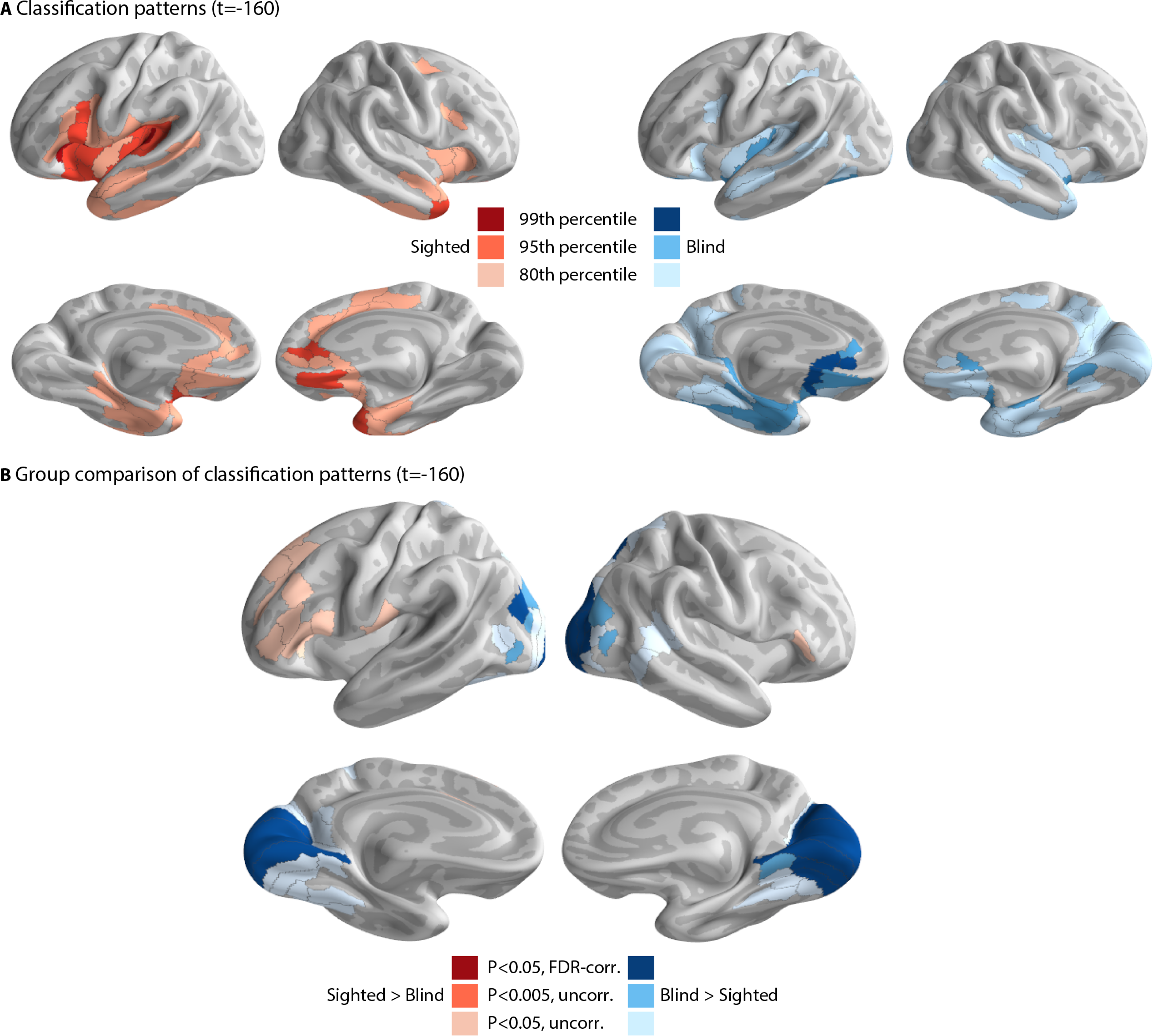
Cortical sources informative for semantic classification. Cortically reconstructed patterns are shown at the time-point of peak classification accuracy in blind participants before button press (−160ms). **(A)** The 99^th^, 95^th^ and 80^th^ percentiles of classification patterns in the sighted (shared of red) and the blind (shades of blue). **(B)** A comparison of the classification patterns in both groups. Results show that the bilateral visual cortex is more informative in blind participants (shades of blue; P<0.05, FDR-corrected, t-test with non-parametric permutations) and the left inferior frontal cortex is more informative in sighted participants (shared of red, P<0.05, uncorrected). The results suggest that systematic group-level features informed classification.

### Cross-subject consistency of neural signatures

We wished to quantify the degree to which the brain responses underlying correct classification were consistent across participants. To this end, in each group separately, we repeatedly trained a classifier on all participants except one, which was used to test the classifier. When averaging all predictions, we found above chance cross-subject generalization in the sighted but not in the blind (p<0.025 at peak time, t-test with non-parametric permutations; Fig. 6). This suggests that, in sighted participants, processing semantic categories is associated with more consistent magnetic field patterns, while the brain responses of blind participants seem to be too variable across participants for cross-subject generalization to succeed. Accordingly, generalization between groups, i.e., training on the all sighted participants and classifying using blind participants, and vice-versa, did not show significant generalization.

**Figure 6.**
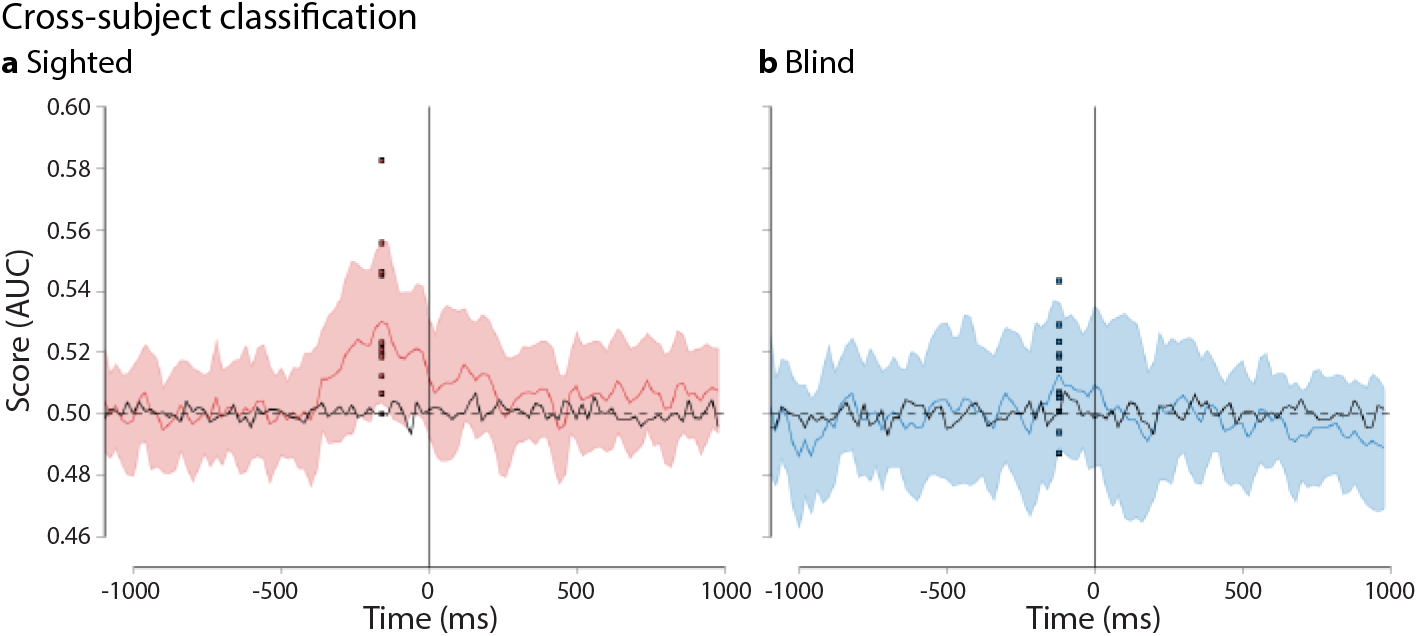
Cross-subject generalization. Classification accuracy across participants in the sighted **(A)** and the blind **(B)**. The X-axis shows time and (ms) where zero corresponds to the button-press and the Y-axis shows the AUC score. The black lines represent decoding accuracy using the dummy models. Shaded areas indicate standard deviation across participants. In each graph, individual values are plotted at the time-point of peak AUC score reflecting classification success of individual participants. The results suggest that patterns related to category discrimination were systematically more consistent in the sighted than the blind participants.

We then tested whether this larger cross-subject variability among blind participants could stem from the known variability in language lateralization in the blind (Lane *et al.*, 2017). Indeed, we found that in the sighted group, there was a consistent left-lateralization in frontal and temporal regions whereas in the blind group, lateralization was highly variable, both in the frontal and in the occipital regions (Supplementary Fig. 4).

## Discussion

### Summary

Enhanced activation versus baseline has been observed repeatedly in the occipital cortex of early blind individuals during a variety of tasks that involve verbal material (see Bedny, 2017 for a review), including tasks requiring access to word meaning (e.g. Noppeney *et al.*, 2003). Those studies however did not establish whether such activations are related to the presence of actual semantic content, or to non-specific task-related processes. In order to address this question, we investigated the spatiotemporal unfolding of access to word meaning in early blind and sighted individuals, using MEG, during a semantic decision task. In blind participants, brain responses to auditory words evolved in the same time window as those observed in sighted participants (Fig. 2A-B). In the blind group only, responses extended to occipital regions (Fig. 2C). Univariate analyses showed that responses in individual participants differed across semantic categories, again in the same time window in both groups (Fig. 3A-B). Crucially, the signal collected in the blind over a cluster of occipital sensors differed across semantic categories, indicating that occipital activations actually carried meaning-related information (Fig. 3C-F). Multivariate analyses confirmed that semantic categories could be discriminated in both groups at the single-trial level in the same time window (Fig. 4), and that the occipital cortex of the blind had a unique contribution to the decoding of word meaning (Fig. 5). Finally, using cross-subject decoding, we also found higher variability in the cerebral implementation of semantic categories in blind than in sighted participants.

The experimental design ensured that category discrimination cannot be attributed to low-level features: Word frequency, number of phonemes, number of syllables, physical duration, and response hand were carefully equated across categories. Moreover, we mitigated inter-item response variability by replicating the analysis while aligning MEG epochs to the button press. Finally, in a control analysis, we checked that differences in response latency could not explain discrimination performance in individual participants, indicating that we actually decoded semantic-related processes and not some correlated feature.

Furthermore, our work stands up to potential criticism put forward by recent epistemological studies, which warn about the divergence of methods based on prediction versus inference (Lo *et al.*, 2015; Bzdok *et al.*, 2018), as well as the risk of misrepresentation when aggregating data across participants (Fisher *et al.*, 2018; Smith and Little, 2018). In our study, multivariate predictive analysis converged with mass-univariate inference that was reproducible across participants (Fig. 3; Fig. 5), and we were able to show that the differences between groups reflect patterns at the individual level (Fig. 3A-C; Supplementary Fig. 1A-C).

### The spatiotemporal unfolding of semantic access

We found that the temporal unfolding of semantic access is quite similar in blind and sighted participants. This is visible in univariate analyses showing discrimination of word categories in individual participants (Fig. 3A; Supplementary Fig. 1A), as well as in multivariate decoding (Fig. 4). In both groups, semantic discrimination overlaps with the usual time window of the N400 component, which is thought to reflect access to word meaning in both the auditory and the visual modalities, based on a huge experimental literature (Kutas and Federmeier, 2011). The broad temporal extent of the N400 is thought to cover stimulus-related activity in the semantic system, with incremental convergence on specific word meaning, modulated by task demands, context and expectations. For instance, in sighted participants, Travis et al. (2013) showed an N400 to auditory words in a semantic matching task, peaking around 400 ms after word onset. Closer to the present task, Chan et al. (2011b) were able to cross-decode semantic category between auditory and visual words, mostly in the 400 to 700 ms window. Earlier discrimination of semantic category may be possible in some cases, particularly using intracerebral recording (Chan *et al.*, 2011a). When considering the spatial domain, the sources of semantic decoding which we observed in both blind and sighted participants (Fig. 5A) also match the known sources of the N400 component, i.e., left-predominant temporal and inferior frontal areas (Lau 2009; Chan 2011; Travis 2013). Only in blind participants, however, there was an additional contribution of the occipital cortex to semantic discrimination, which is compatible with EEG studies showing a less frontal topography of the N400 in the blind than in the sighted (Röder *et al.*, 2000; Glyn *et al.*, 2015). Importantly, this discrimination unfolds in the same N400 period as in frontotemporal areas (Fig. 3B; Fig.5B), suggesting that occipital activity contributed to actual semantic access and not only to post-decisional processes.

Why did the occipital contribution to word semantics in the blind escape previous studies? As discussed before, some studies have shown general occipital sensitivity to semantic processing without trying to discriminate between semantic categories (Amedi *et al.*, 2004; Bedny *et al.*, 2011). Other fMRI studies, using univariate methods, found differences between word categories in temporal cortex but not in occipital regions (Noppeney *et al.*, 2003; Struiksma *et al.*, 2011; Bedny *et al.*, 2012), or even focused the analyses only on the ventral occipitotemporal cortex (e.g. Mahon *et al.*, 2009). However, two previous studies should be considered as immediate context to the present work. First, Schepers et al. (2012) used MEG to show an effect of semantic congruency in V1 of blind participants, in the N400 time window, using haptic and auditory presentation of objects. This study showed that semantic processes may indeed be detected in the occipital lobes of blind participants with MEG, but they did neither use verbal material nor test category discrimination. Second, van den Hurk et al., (2017), also using non-verbal stimuli (audio clips), showed with fMRI that decoding category membership was possible in V1 in the blind. The present study fills the critical gap of demonstrating the contribution of the occipital lobes to word meaning in the blind, in the same time window as conventional semantic processes.

### Variability as a consequence of neuronal plasticity?

We found that the occipital cortex of the blind can discriminate semantic categories. But we also found that the signals underlying this discrimination do not generalize across subjects. In contrast, cross-subject generalization was more successful in sighted participants. Blind participants may differ among them in the contribution of various anatomical regions to semantic coding, but also in the precise small-scale configuration of neural activity within those regions. In support of processing in different regions, we found that classification patterns in blind participants showed inconsistent hemispheric lateralization, whereas in sighted participants there was a consistent left-lateralization in the inferior frontal cortex (Supplementary Fig. 4). This is in agreement with previous evidence that language areas are less left-lateralized in three different groups of blind participants when compared to sighted participants (Lane *et al.*, 2017). Therefore, the less consistent magnetic field patterns indicated by the lack of generalization across blind participants could result from this lack of consistent lateralization. Independently, the configuration of neuronal sources at similar macroscopic locations could also differ (e.g. dipoles with the same center but different orientation and polarity), giving rise to distinct dipolar field patterns that would also impair cross-subject generalization. Both accounts are not exclusive, and suggest that plasticity could follow unique patterns across blind individuals. In this context, computational micro-circuit modelling (Jones *et al.*, 2009; Khan *et al.*, 2015; Sherman *et al.*, 2016) may help further elucidate the anatomical scales of plasticity and its impact on MEG and EEG observables when applied of the visual cortex in blind participants.

### Conclusion

Discrimination between different semantic categories showed similar temporal unfolding in blind and sighted participants. In the blind only, this process involved the occipital cortex in addition to frontotemporal regions. Moreover, it seems that the neural implementation of semantic processing is more variable across blind participants than across sighted participants.

Further work is needed to elucidate the exact role of the occipital cortex in semantic processing. Using other tasks, it may be possible to dissociate between a role in the storage of semantic information or in the executive processes manipulating semantic information, as suggested by the controlled semantic cognition model (Ralph *et al.*, 2017). Importantly, the inter-subject variability we found should be replicated by future studies explicitly designed to uncover its sources and implications.

## Acknowledgments

The authors would like to thank Nathalie George and Denis Schwartz for their help in MEG acquisition, Celine Cluzeau for coordinating subject recruitment and data acquisition, Alexandre Gramfort, Benjamin De Haas, Guillaume Dumas, Jacobo Sitt, Marijn Van Vliet (in alphabetical order), the Parietal Team and the PICNIC Lab for stimulating discussions and feedback on our work.

