## Supplementary materials for "Semantic coding in the occipital cortex of early blind individuals"

### Supplementary materials and methods

#### Experimental design and stimuli

Across semantic categories and across phonemic categories, words did not differ in terms of number of phonemes (both $F(2, 81) < 1$, $P=1$, ANOVA), of frequency of occurrence (both), and of number of syllables ($F(2, 81)=0.35$, $P=0.70$, $F(2, 81)=2.17$, $P=0.12$). Stimuli and instructions were synthetically generated using the Text-to-Speech function built in OSX 10.9.5 (Apple Inc., CA, United States). Sound files were normalized (maximum amplitude set to -1.0dB) using Audacity (version 2.1.2, http://audacity.sourceforge.net/). Sound file duration did not differ across the three semantic categories ($F(2, 87)=1.04$, $P=0.358$; Tukey's test for the three pairs: $P=0.340$, $P=0.600$, $P=0.897$)

#### MEG source localization

To estimate cortical neuronal dynamics from the observed sensor array time series we approximated a numeric solution to the biomagnetic inverse problem using cortically constrained Minimum Norm Estimates (MNE) with l2 regularization. The MNE solver yields a linear spatial filter that projects MEG data onto a predefined grid of cortical source locations. Concretely, we computed the MNE solution by fitting the regularized linear regression model to the MEG data based on the forward solution, which quantifies profiles of magnetic field propagations from source locations to MEG sensors from the individual anatomy and conductivity models. As MEG sources primarily reflect population synchrony of big pyramidal layer 4 neurons (Hämäläinen et al. 1993) we placed all source locations on the cortical sheet, hence the cortical constraint prior. We used a single-layer boundary element model (Hamalainen and Sarvas 1989) constrained by the individual anatomical MRI and extracted the cortical surfaces with FreeSurfer with subsampling to about 4098 vertices per hemisphere yielding a surface area of about $24mm^2$ for each source. To minimize violations of the model assumptions of multivariate Gaussian noise, spatial whitening is commonly applied to the MEG data and the forward model based on the noise covariance matrix (Engemann and Gramfort 2015). We estimated the noise covariance from 200ms of baseline segments prior to the stimulus onset using an optimized shrinkage estimator (Engemann and Gramfort 2015). To obtain the final spatial filter we applied depth-weighting (gamma = 0.8) and the loose orientation constraint (loose = 0.2; Lin et al. 2006), both default settings in MNE-Python. This constraint was a compromise between pooling the current orientations and relying on the correctness of the individual curvature information when computing the sign of the signal by weighting the source variances of three dipole components that define the tangent space of the cortical surfaces. We then set the regularization parameter controlling the *lambda2* following the MNE software's standard practice of dividing 1 by the squared signal-to-noise ratio ($1/SNR^2$), where the SNR of event-related data is conventionally assumed to be 3. To make solutions more comparable across subjects, we applied dynamical statistical parametric mapping (dSPM) noise normalization (Dale et al. 2000) which recasts the estimated source current amplitudes relative to its noise-floor, yielding a pseudo F-statistic. We, moreover, used the FreeSurfer routines described above to warp the dipole grids from each individual to the FreeSurfer average brain.

As the MNE solutions is linear and consists in a single matrix multiplication it can be applied to either the time-series, complex-valued Fourier coefficients or other linear transforms under conditions of identical noise. Here we made use of this property to estimate the sources associated with our linear decoding patterns.

To alleviate the multiple comparison problem, prior to performing group-level statistics, we summarized our inverse solution using the Human Connectome Project (HCP) cortical parcelation (Glasser et al. 2016) with 360 functionally-defined regions of interest (ROI) covering the entire cortical surface. Concretely, we averaged the dSPM values for each subject in each ROI.

#### Machine Learning

We used a multivariate pattern classification approach for decoding semantic category. Accordingly, we trained a linear pattern classifier to approximate a function that maps MEG signals to condition pairs. To estimate out-of-sample performance we employed a seven-fold group cross-validation where grouping ensured that words used during training were never used during testing. Seven was the largest number of folds possible given this grouping-constraint. We used an l2-regularized linear model with sigmoid loss function (Logistic Regression) with a constant regularization parameter of C = 1 and trained it on all magnetometer and gradiometer sensors without feature selection. Trials were dropped when necessary for equating the number of right and left presses across categories to prevent bias. We considered single sensor topographies time-point by time-point (e.g. King et al. 2014) and assessed performance using the area under the curve (AUC) of the Receiver-Operator-Characteristic. To compute cross-subject generalization, we used leave-one-subject-out cross-validation. Binary decoding was performed on the three category pairs whose results were then averaged. All machine learning was performed using the scikit-learn software (Pedregosa et al. 2011).

To facilitate interpretation of the learned model parameters, which, due to their conditional nature, may reflect either noise or signal, we marginalized the parameters by multiplication with the feature covariance (Haufe et al. 2014). This resulted in the classification patterns for each category pair. We then estimated the sources underlying the patterns by applying the linear MNE inverse operator. To satisfy the empirically estimated noise model obtained from covariance estimation and preserve correct scaling of the inverse solution, we summed the pattern and the evoked response prior to source localization. We then reconstructed the pattern of each category pair by subtracting the corresponding source localized evoked response. We then considered absolute values, as the sign of this contrast should be driven by individual source geometry which would not add up across subjects. As a result, we obtained a cortical map indicating the strength of the contributions to either negative or positive model coefficients. Finally, in order to assess the lateralization of the classification patterns in source space, for each subject, we first subtracted the right hemisphere from the left hemisphere and vice versa. Then, for each ROI in source-space, we computed the percentage of the subjects from each group where the value of the difference was above 0.5.

### Supplementary results

#### Category discrimination aligned on button press

Previous analyses showed that semantic discrimination in both groups, as well as differences in discrimination between groups, cover roughly late components (i.e. discrimination after 400 ms and group differences after 600 ms). This may suggest that aligning epochs on word onset may not be optimal, as it maximizes the between-word variability in the time point at which a category decision is reachable. Hence, synchronizing on word onset may result in an overlap between data from evidence accumulation prior to decision (in words with a late decision point), and data from response-related or later processes (in words with an early decision point). Alternatively, aligning epochs on the button press may help averaging out such between-word variability by amplifying signals coupled with the decision process. We therefore repeated the discrimination analysis after aligning trials to button press. This yielded similar results to word onset alignment. Individual category discrimination was found in all but two sighted participants (Supplementary Fig. 1A-C), and significant group difference in discrimination was found in a cluster encompassing posterior sensors between -460 and -20 ms ($P<0.003$;

Supplementary Fig. 1D-E). The discrimination index was higher in blind than in sighted participants in the occipital cortex, and the difference peaked at -120 ms relative to button press (Supplementary Fig. 1F). The cluster with significant between-group differences in category discrimination had a longer duration when aligning to button-press (440 ms) than when aligning to word onset (320 ms). As aligning trials on button press seemed to increase the power of the analysis, we proceeded with this option in the analyses that follow.

### Supplementary Figures

#### Supplementary Figure 1

**Neural correlates of semantic category discrimination relative to button press.** Panels A-C depict individual analyses. **(A)** The X-axis shows time (ms) where zero corresponds to the button press and each row on the Y-axis shows the result of one subject, sighted in red and blind in blue. Each bar represents a significant spatiotemporal cluster using the magnetometer sensors, bar width is inversely proportional to the p-value and cluster overlap renders colors darker. At least one significant cluster close to the button press can be seen in all but two sighted participants. **(B)** Temporal overlap of clusters in Sighted (red) and blind (blue) participants. Colors represent the percentage of participants with a significant cluster that includes a given time point. The lightest color represents time points where less than 50% of the participants had a significant cluster. **(C)** Spatial overlap of clusters, showing the percentage of participants with a significant cluster that includes a given sensor. The color scheme follows the description in panel B. Panels D-F depict comparisons between groups. **(D)** The X-axis shows time (ms) where zero corresponds to the button press and the Y-axis to the average category discrimination in sighted (red) and blind (blue) participants. The yellow bar indicates the temporal extent of the spatiotemporal cluster with significant difference across the groups (P=0.003, spatiotemporal permutation clustering). **(E)** The average topography of the group difference in semantic discrimination during the time of the significant cluster (from -460 to -20 ms). Sensors appearing in the cluster are highlighted in white. The peak difference between sighted and blind participants (in the time-window of the cluster in panel D) when comparing the average of all pair-wise contrasts. It shows a stronger effect in the occipital cortex of blind participants and no regions with a significant effect in sighted participants. The results suggest that occipital sensors were most sensitive to category discrimination in the blind as compared to the sighted participants and point at specific contributions from the visual cortex.

#### Supplementary Figure 2

**Cortical sources informative for semantic classification at different time points.** Similar to Figure 5. Cortically reconstructed patterns informative for semantic classification at the time-point of peak classification accuracy in the sighted before button press (-80 ms). **(A)** The $99^{th}$, $95^{th}$ and $80^{th}$ percentiles of classification patterns in the sighted (shared of red) and the blind (shades of blue). **(B)** A comparison of the classification patterns in both groups. Results show that the bilateral visual cortex is more informative in blind participants (shades of blue; $P<0.05$, FDR-corrected, t-test with non-parametric permutations) and the left inferior frontal cortex is more informative in sighted participants (shared of red, $P<0.05$, uncorrected).

#### Supplementary Figure 3

**Cortical sources informative for semantic classification at different time points.** Similar to Figure 5. Cortically reconstructed patterns informative for semantic classification at the time-point of peak classification accuracy in the sighted and blind participants after button press (+60, and +80 ms, respectively). **(A-B)** The $99^{th}$, $95^{th}$ and $80^{th}$ percentiles of classification patterns in the sighted (shared of red) and the blind (shades of blue). **(C-D)** A comparison of the classification patterns in both groups.

Results show that the bilateral visual cortex is more informative in the blind participants (shades of blue; $P<0.05$, FDR-corrected, t-test with non-parametric permutations) and the left inferior frontal cortex is more informative in the sighted participants (shared of red, $P<0.05$, uncorrected).

##### Supplementary Figure 4

Inter-subject variability at peak cross-subject classification (-160 ms for the sighted and -120 ms for the blind). Left hemisphere maps show the percentage of participants with a stronger contribution from the left than the right hemisphere to classification, and conversely for right hemisphere maps. A high overlap in a region reflects its consistent lateralization across participants. In the sighted, there is a consistent left-lateralization in frontal and temporal regions. In the blind participants, lateralization is highly variable, both in the frontal and in the occipital regions.

#### Supplementary Tables

##### Supplementary Table 1

The list of nouns using in the semantic decision task.

| **Animals** | | | **Manmade** | | | **Plants** | | |
|---|---|---|---|---|---|---|---|---|
| abeille | merle | saumon | marmite | fléchette | clochette | mâche | herbe | pomme |
| baleine | mésange | goéland | manège | épingle | torchon | papaye | cresson | origan |
| carpe | zèbre | dauphin | armure | écharpe | bobine | carotte | trèfle | laurier |
| crapaud | bécasse | fauvette | gare | chaise | orgue | lavande | persil | pommier |
| chameau | chèvre | faucon | barrette | cerceau | chaudron | pastèque | verveine | romarin |
| sardine | reptile | mollusque | balai | flèche | brosse | sapin | fraise | orange |
| chamois | blaireau | vautour | cadran | veston | lotion | cannelle | sésame | roquette |
| chacal | lézard | tortue | chalet | évier | sofa | navet | laitue | olive |
| hanneton | guépard | brochet | chaloupe | béquille | totem | chardon | mélèze | rosier |
| vache | aigle | coq | cahier | béton | broche | maïs | raisin | oseille |

Supplementary Figure 1

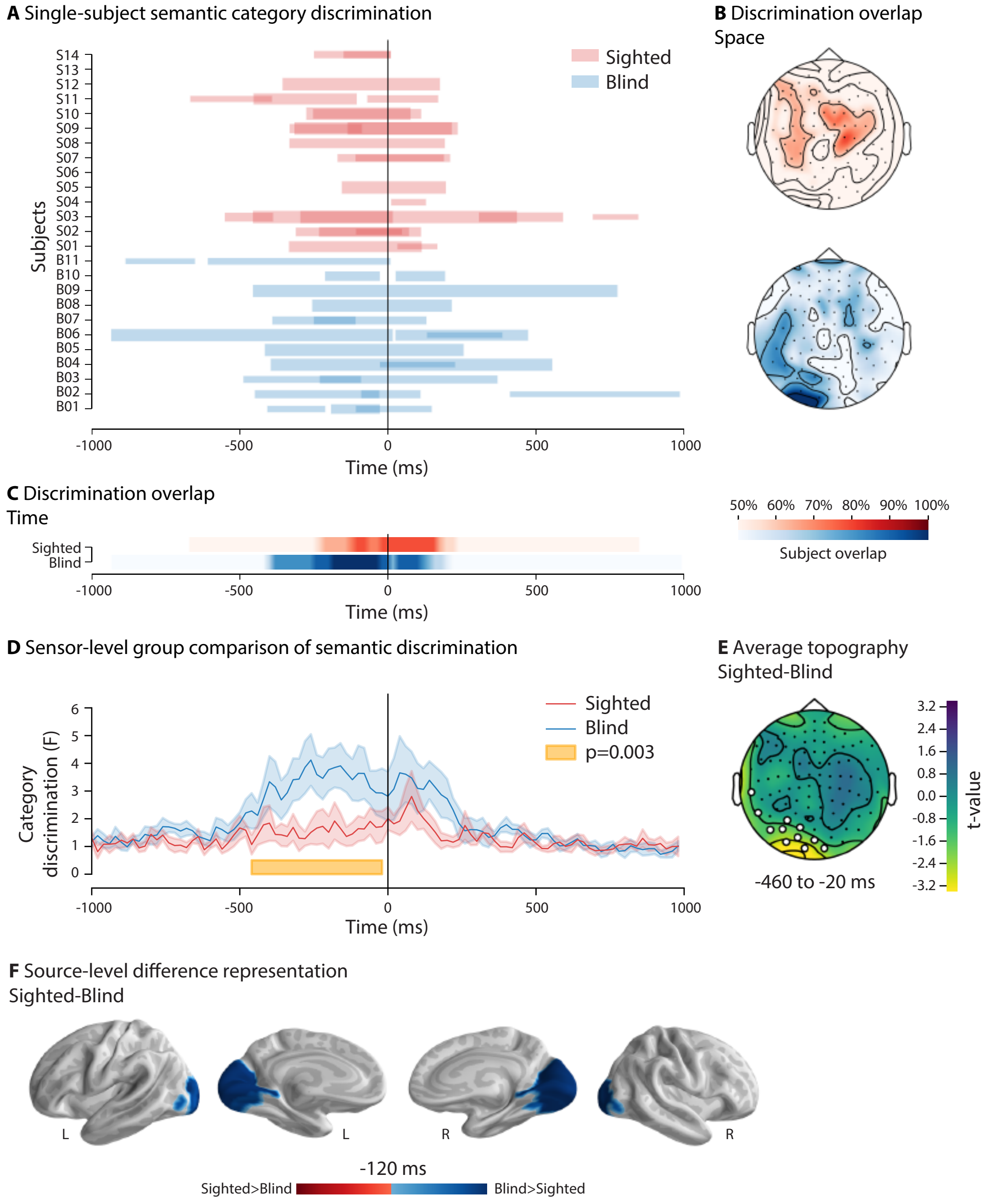

Supplementary Figure 2

**A** Classification patterns (t=-80)

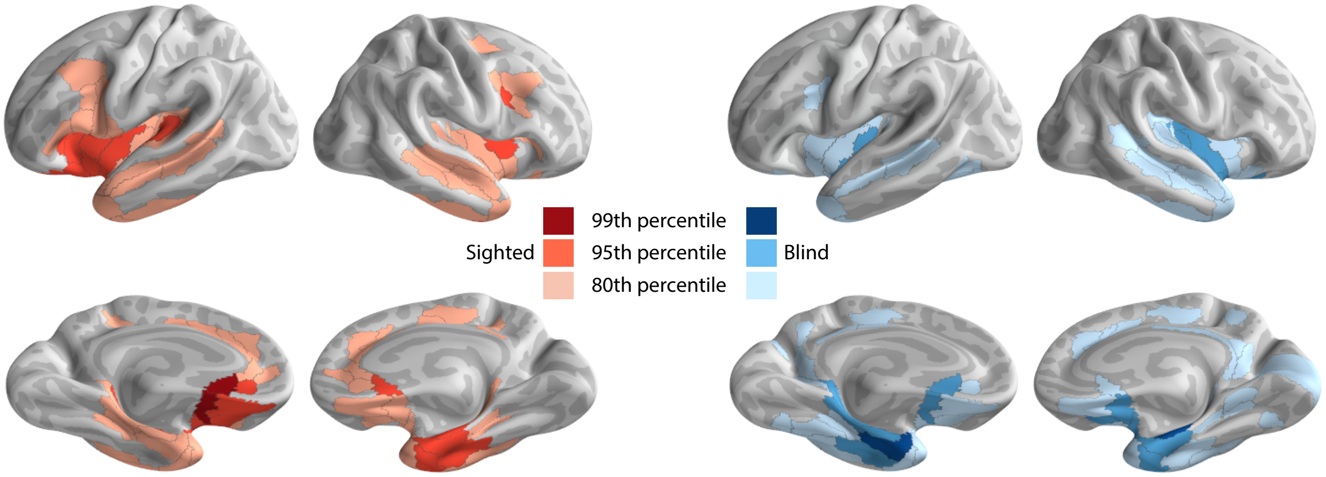


**B** Group comparison of classification patterns (t=-80)

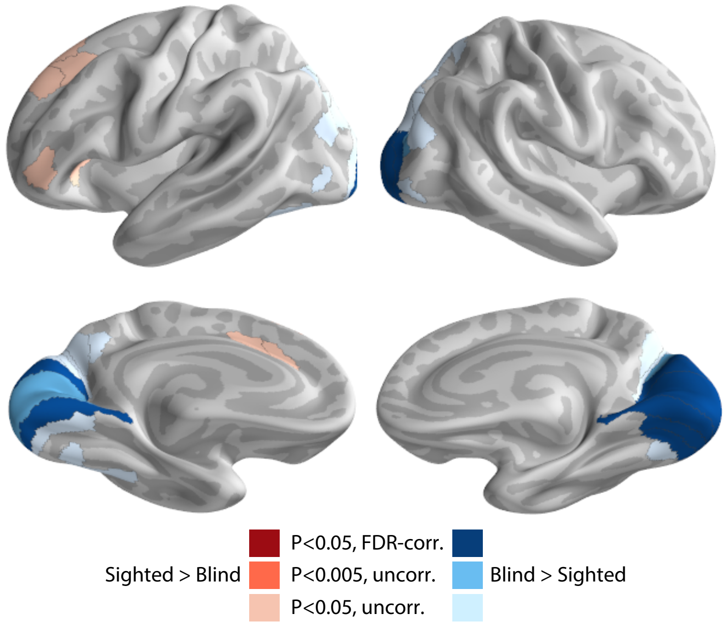

Supplementary Figure 3

**A** Classification patterns (t=60ms)

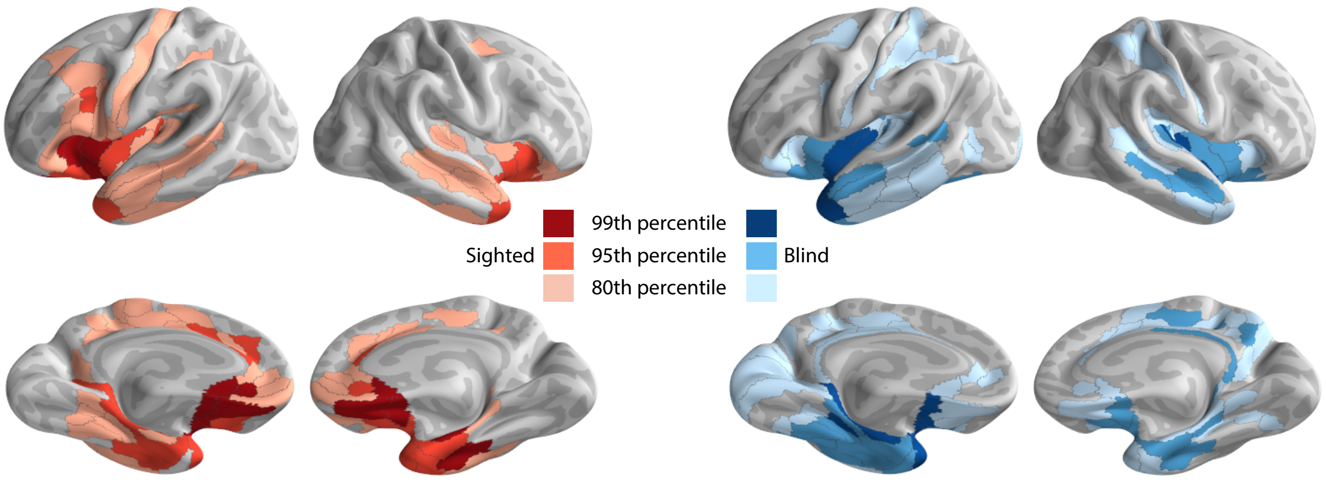


**B** Classification patterns (t=80ms)

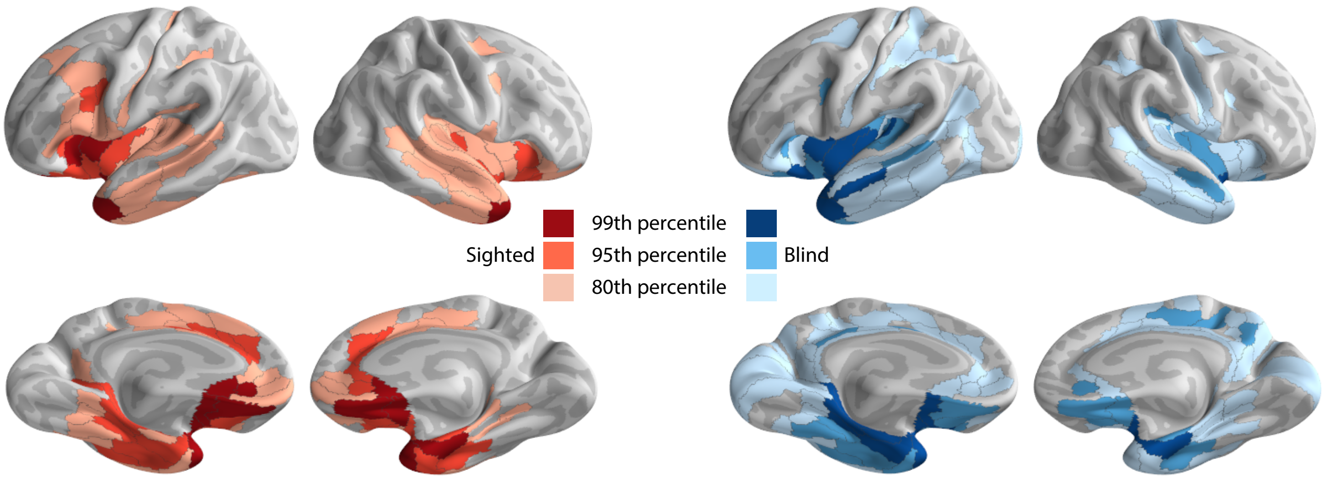


**C** Group comparison of classification patterns (t=60ms)

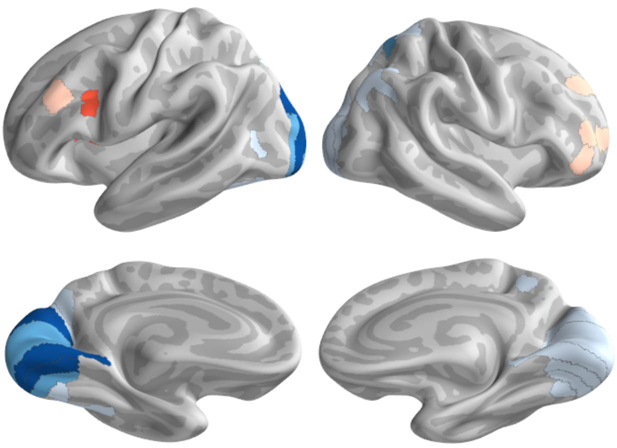

**D** Group comparison of classification patterns (t=80ms)

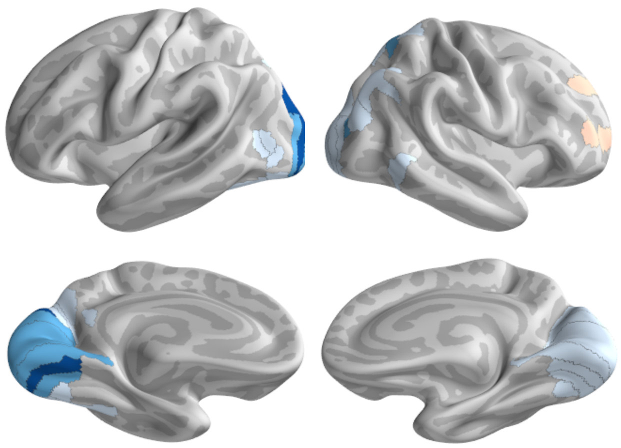

Sighted > Blind
P<0.05, FDR-corr.
P<0.005, uncorr.
P<0.05, uncorr.
Blind > Sighted

Supplementary Figure 4

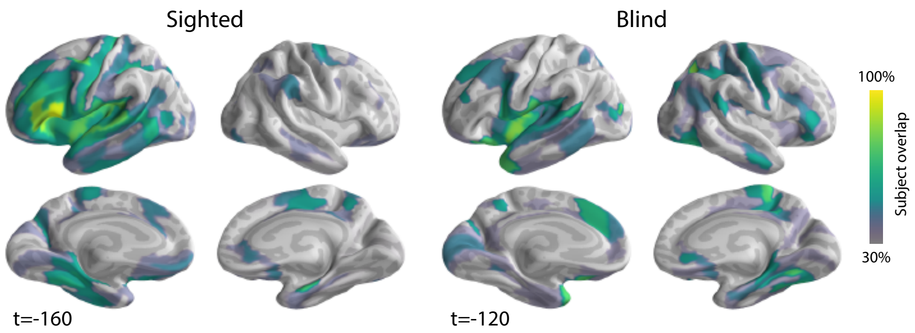
